# Phyla-specific Correlated Dynamics in Tropomyosin

**DOI:** 10.1101/301028

**Authors:** J.K. James, V. Nanda

## Abstract

Tropomyosin (Tpm) is a continuous α-helical coiled-coil homodimer that regulates actinomyosin interactions in muscle. We examined extended molecular simulations of four Tpms, two from the vertebrate phylum Chordata (rat and pig), and two from the invertebrate Arthropoda (shrimp and lobster), and found that despite extensive sequence and structural homology across metazoans, dynamic behavior – particularly long range structural fluctuations – were clearly distinct between phyla. Vertebrate Tpms were flexible and sampled complex, multi-state conformational landscapes. Invertebrate Tpms were rigid, sampling highly constrained harmonic landscapes. Filtering of trajectories by PCA into essential subspaces showed significant overlap within but not between phyla. In vertebrate Tpms, hinge-regions decoupled long-range inter-helical motions and suggested distinct domains. In contrast, crustacean Tpms lacked significant long range dynamic correlations – behaving more like a single rigid rod. Although Tpm sequence and structure has highly conserved over the last 0.6-billion years since the split of ancestral bilateria into protostomes and deuterostomes, divergence seems to have occurred at the level of long-range correlated dynamics, reflecting adaptations to phyla-specific requirements of actin binding and muscle contraction.

## INTRODUCTION

Tropomyosin (Tpm) is an ancient protein, found throughout diverse metazoan phyla in cytoskeleton and muscle thin filaments as an actin binding protein (Fig. 1) (3). It molecular structure is a parallel, extended coiled-coil homodimer that is almost always 281-284 residues long. A number of conserved sequence features are found throughout the animal kingdom: periodic clusters of core-facing alanine residues can induce local instabilities that modulate global flexibility and function (4), and periodic surface-facing acidic residues that form actin binding sites. These and other Tpm sequence/structure features are practically universal across higher eukaryotes (3, 5).

**Figure 1.**
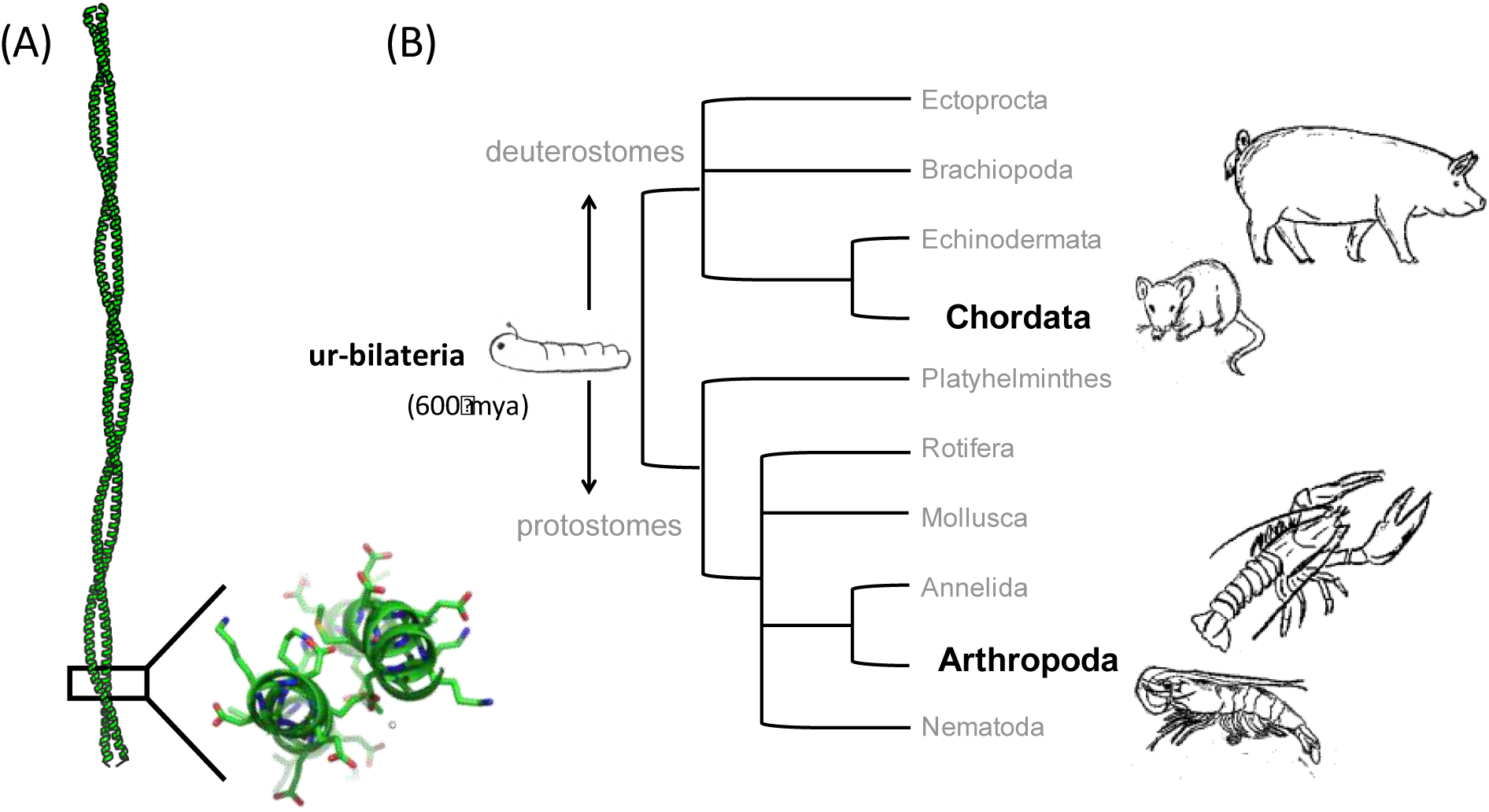
Evolutionary tree of Bilateria. **(A)** Tpm forms an extended parallel α-helical coiled-coil dimer whose sequence and structure is highly conserved over hundreds of millions of years of metazoan evolution. **(B)** We examine the structure and dynamics of Tpm from two phyla: Chordata (i.e. vertebrates) including rat and pig, and Arthropoda, focusing on shrimp and lobster from the Crustacea sub-phylum. Chordates and arthropods belong to two major branches of Bilateria (organisms with bilateral symmetry), the protostomes and deuterostomes. The protostome/deuterostome split from an ancient ur-bilateria is proposed to have occurred over 600 million years ago (1, 2).

Despite this high degree of conservation across metazoans, there are features of Tpm that present in a phyla-specific manner. Arthropod Tpms are pan-allergens, giving rise to food and respiratory allergies (6). Vertebrate Tpms, on the other hand, generally do not cause allergy – although recent exceptions have been noted in the clinical literature (7). Furthermore, patients with Tpm-based allergies to foods such as shellfish do not report cross-sensitivities to foods such as pork or chicken, which contain significant amounts of vertebrate Tpm. In a sense, the phyletic separation of allergens and non-allergens is the epidemiological equivalent of classic pre-gene sequencing ‘immunological distance’ experiments, where the extent to which antibodies raised against sera from one organism react with another was used as a proxy for phylogenetic distance (8, 9). With extensive modern genomic and protein structure information, evolutionary relationships can be determined with significantly improved precision. Even at the molecular level, the sequence identity between arthropod and vertebrate Tpms is 50-60% and structure prediction tools indicate both species form coiled-coil topologies (**Fig. S1**). Analysis of sequence and structure alone do not effectively discriminate allergenic and non-allergenic forms of Tpm (10).

Vertebrate and crustacean Tpms exhibit different dynamics in molecular simulations. In muscle contraction, Tpm adopts a curved conformation, allowing it to wrap around an actin filament. Distinct models have been proposed for how this curvature is achieved: a gestalt binding model assumes a rigid, curved conformation that complements the actin filament (11), and a hinge model where destabilizing residues, primarily within the core, confer flexibility to allow Tpm to sample conformations for optimal binding (12). The gestalt binding model is supported by observed curvature and persistence length several times the length the coiled-coil as seen by electron microscopy (EM), molecular dynamic simulations (MD) (13, 14), and cryo-EM studies of conformational transitions of Tpm along actin (15). In support of the hinge model, non-canonical core packing has observed in both high resolution crystal or NMR structures produce local flexibility (4, 16, 17). Biophysical studies of unfolding transitions suggest cryptic structural domains whose determinants are not obvious (18, 19), but may contribute to local bending of the coiled-coil.

We explore whether molecular evolution has selected for Tpm dynamics behavior over sequence determinants of binding and large scale modifications to structure. Despite its long coiled-coil structure, Tpm is sensitive to single point mutations, which can significantly alter actin binding (12, 20, 21). Additionally, short time MD simulations demonstrate a pattern of local flexibility over destabilizing regions in the Ala clusters in the core (22). Minor changes in sequence can have dramatic impacts on the stability and cooperativity of Tpm folding domains, impacting actin binding and actinomyosin ATPase activity (20, 21, 23).

Comparative studies of protein dynamics from distant homologs may provide new insight into the relationship between sequence, structure, dynamics and function. Much of Tpm structure and function have been studied in the context of the vertebrate skeletal muscle system. Similar studies at the molecular level are lacking for tropomyosin and its role in invertebrate muscle contraction. Actinomyosin regulation in invertebrates occur by a dual control mechanism including both the typical actin-controlled mechanism with Ca^2+^ activation of the troponin-tropomyosin-actin complex and a myosin-controlled mechanism involving direct Ca^2+^ binding to myosin (24, 25). The evolutionary consequences of these mechanisms on Tpm dynamics are largely unknown.

## RESULTS AND DISCUSSION

### Analysis of Global Dynamics

We used MD simulations to study the collective dynamics of Tpm. All-atom explicit solvent MD simulations for mammalian (rat and shrimp) and crustacean (shrimp and lobster) Tpms were performed for 160 ns using the AMBER simulation platform (26). Starting structures were derived from the low-resolution full-length crystal structure of pig Tpm – PDB ID 1C1G (27). Rat, shrimp and lobster sequence were threaded onto this structure and subject to sidechain optimization in protCAD (28, 29).

Two global order parameters: root mean squared deviation (RMSD) and radius of gyration, show the distinct dynamics between the two phyla examined. Both parameters are sensitive toward global motions, and the latter is sensitive to the curvature of the coiled-coil. Distinct patterns of dynamics are observed between the two phyla (Fig. 2). Rat and pig Tpm show an overall curved conformation that resembles previous simulations of the Lorenz-Holmes Tpm model derived from vertebrate sequences (13). Its shows an anharmonic distribution suggesting a rugged landscape with multiple conformations explored. In sharp contrast, the average conformation in shrimp Tpm is linear, and its conformational distribution is tightly Gaussian with minimal deviations from the average structure.

**Figure 2.**
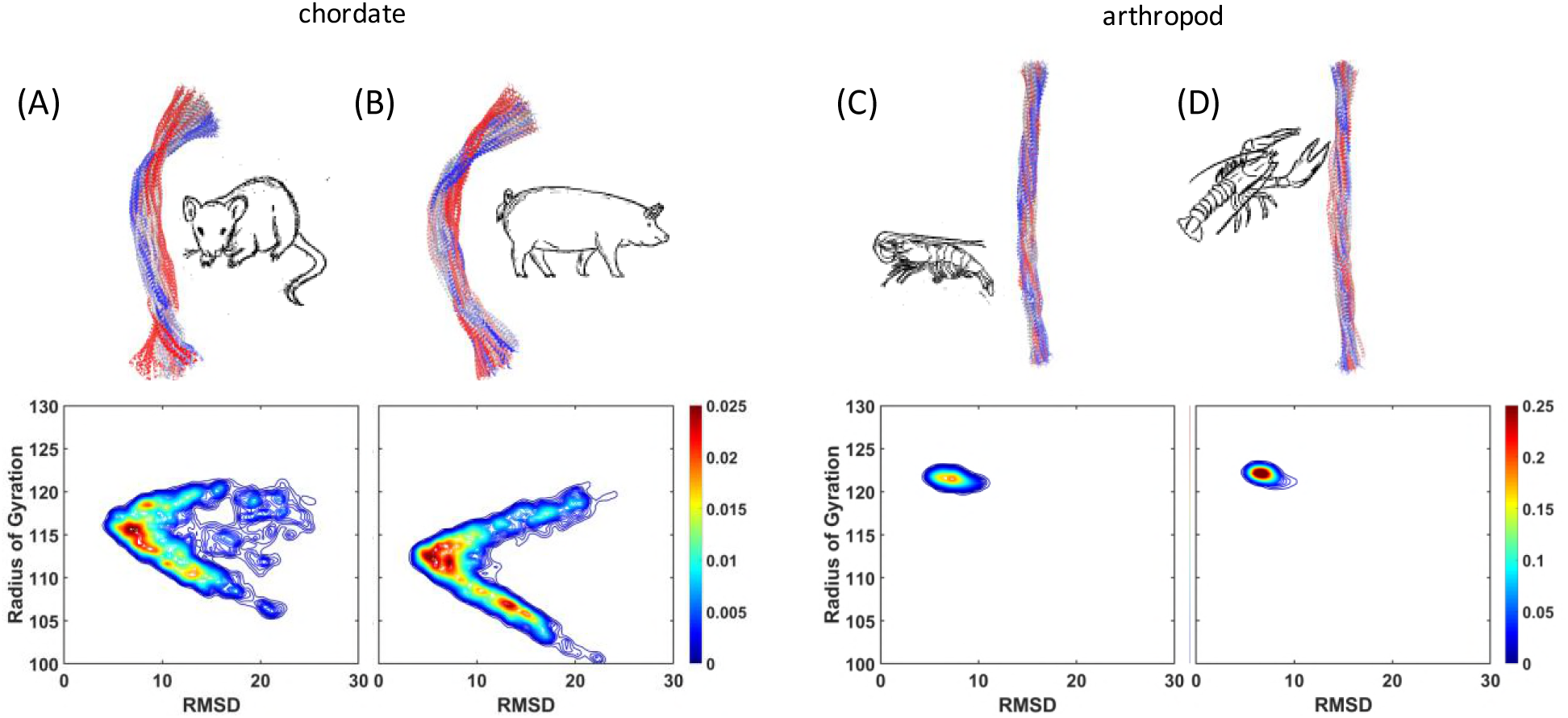
Simulated Conformational Space. Snapshots of structures along an MD trajectory are overlaid for **(A)** rat, **(B)** pig, **(C)** shrimp and **(D)** lobster Tpm. Plotted below are probability density contour plots of RMSD from the average structure versus radius of gyration.

Principal component analysis (PCA) was used to filtering large scale dynamics into essential subspaces (30, 31). The first principal component is an indicator for global anharmonic dynamics associated with transitions between multiple conformational states (32, 33). Anharmonic modes in general represent the minority of principal components but represent the majority of mean square fluctuations (MSF) observed in the dynamics; consistent with the fact that collective dynamics across conformational states generates large scale deviations in structure. Fluctuations of Cα coordinates with respect to the mean position were used for to generate a covariance matrix, which is then diagonalized to generate principal components of the system. Distributions of projecting Cα fluctuations from the trajectory along the top principal components are calculated along with MSF from their associated eigenvalues (Figs. 3, **S2**). In rat and pig Tpms, projections along the first principal component yielded an anharmnonic distribution accounting over half the total mean square fluctuation (Figs. 3A, 3C, **S2A, S2C**). Projections along the second principal component were significantly more harmonic, but associated with a more than three-fold smaller MSF. Projections along the first three principal components in shrimp and lobster Tpm produce a nearly Gaussian distribution, with no mode dominating MSF as in rat (Figs. 3B, 3D, **S2B, S2D**). Distributions of projections were narrow reflecting smaller global motions. The top two modes were also nearly Gaussian and appeared as mirror images in their distribution. The similarity in their distribution is reflected in their equivalent MSF, suggesting an artificial separation of isotropic dynamics.

**Figure 3.**
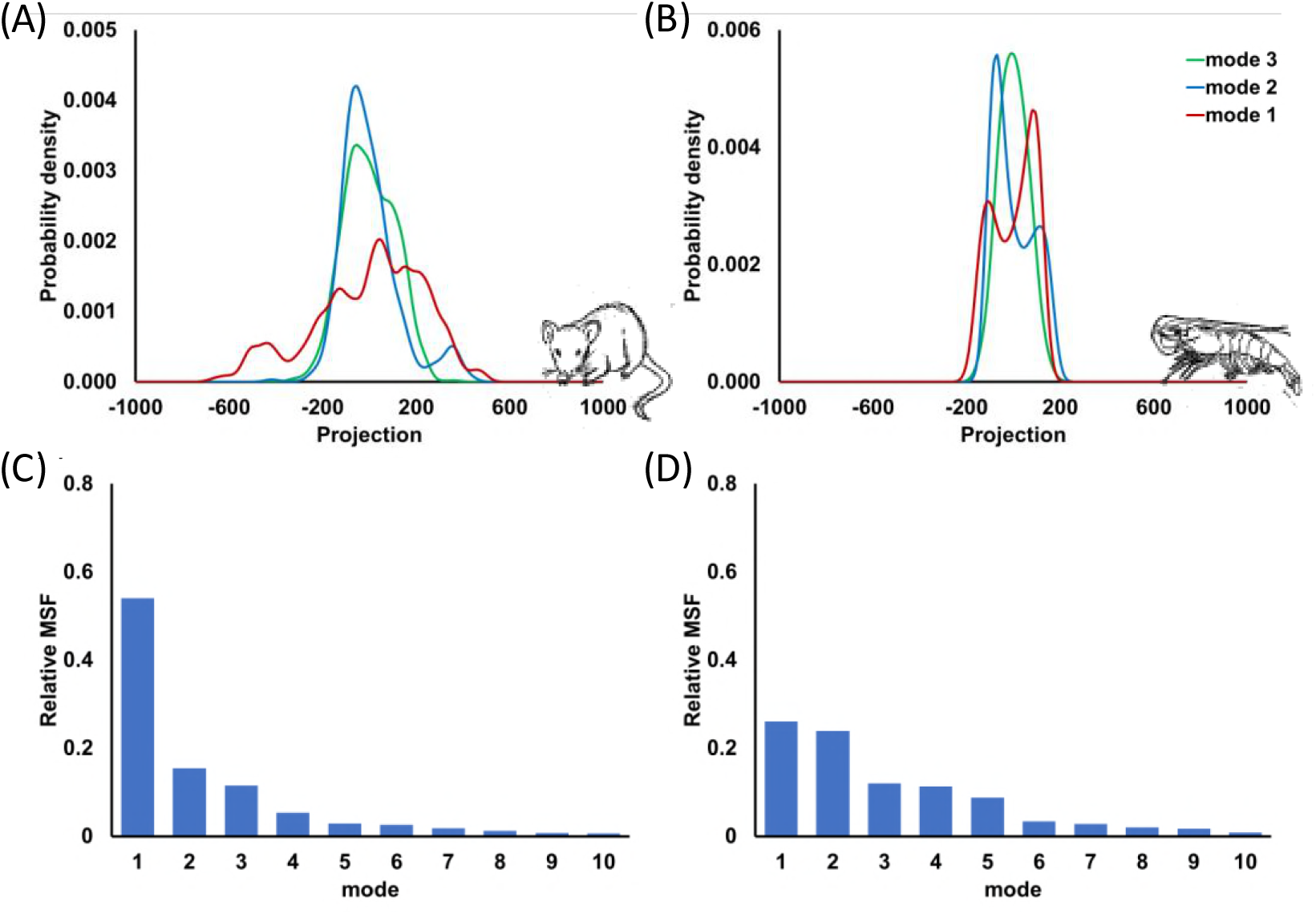
Fluctuations along top Principle Components. Probability Densities along top three principle components calculated directly from MD trajectory of (A) rat and (B) shrimp Tpm. In addition, mean square fluctuation as a ratio to total fluctuation is calculated for top 10 modes in simulations of **(A)** rat and **(B)** shrimp Tpm.

Two basic modes of global dynamics in Tpm were observed: a curved conformation generated from a rugged conformational landscape, and a straight conformation in a single Gaussian well. When comparing the first principal component across all simulations, vertebrate and crustacean Tpms shared over 70% subspace overlap within phyla, and little subspace overlap between phyla (**Table S1**).

### Dynamic Domains

Large shifts in global order parameters such as RMSD or radius of gyration seen in mammalian Tpm require correlated fluctuations along the backbone (34). Correlated dynamics could reveal collective motions of residues that create dynamic domains within the continuous coiled-coil. Domains might be identified using covariance matrix, where the magnitude of covariance corresponds to degree of correlation between residue fluctuations (35). Given the linear nature of the coiled-coil structure, the sign of the correlations matters less than then magnitude. Therefore, the covariance matrix can be represented as the root squared sum of covariance across all three Cartesian coordinates. Additionally, the covariance matrix allows for a hierarchal visualization of intra- and inter-domain correlations. As an example, within a three-domain rigid body system, correlations extending off-diagonal correspond to domain regions, whereas correlations beyond the diagonal show domain-domain associations (Fig. 4A).

**Figure 4.**
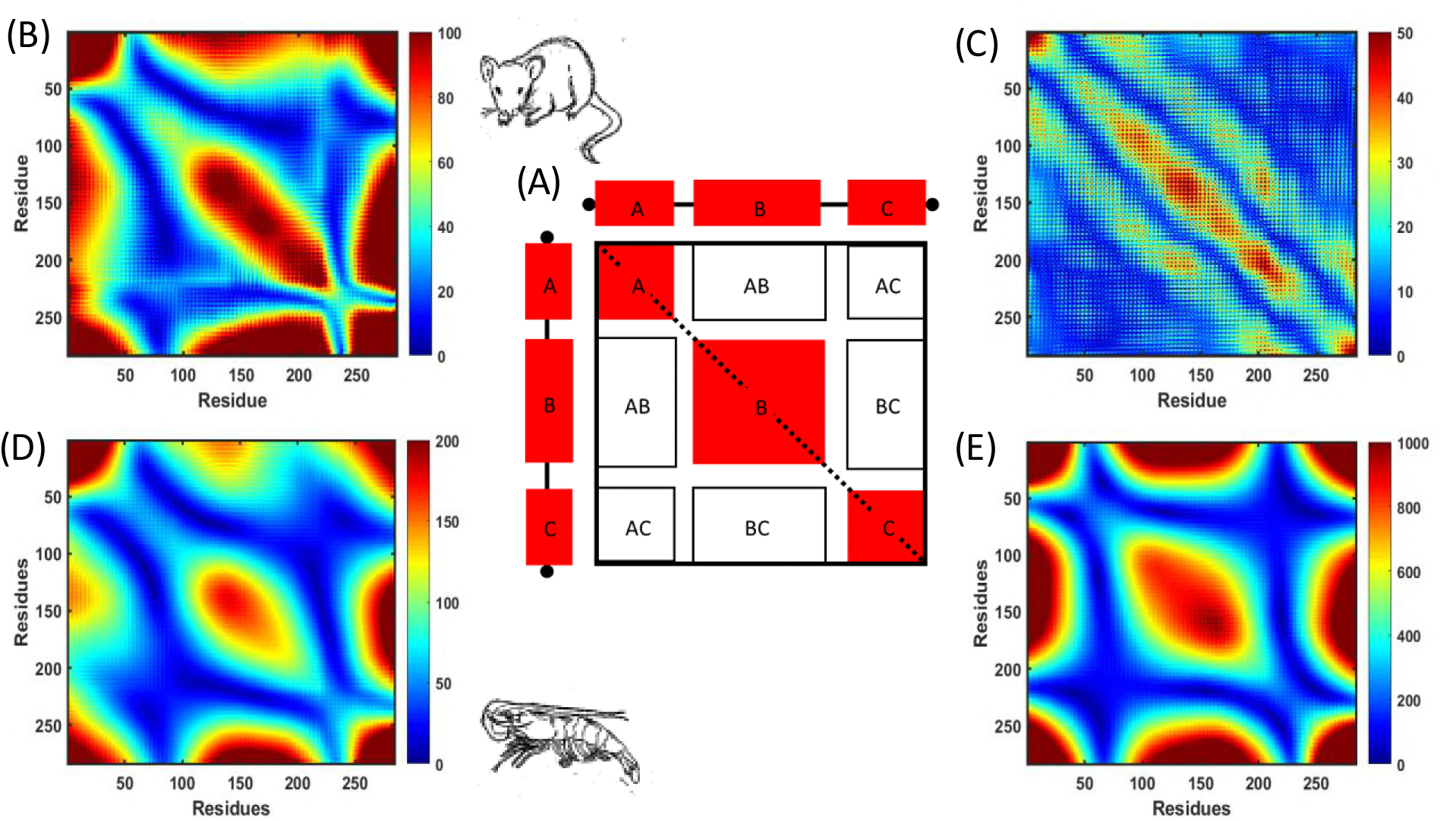
Domain mapping with covariance of residue fluctuations. A covariance matrix was calculated using atomic fluctuations of Cα. (A) A schematic of a covariance matrix in an idealized three rigid-body domain structure, where the three off-diagonal domains are labeled as red blocks with domain-domain correlations outlined in black. Covariance map calculated from MD simulations as the root square sum of cross-correlations produced from Cα fluctuations in X, Y and Z directions for (B) rat and (C) shrimp Tpm. Similarly, a covariance matrix was calculated based on normal modes derived from an anisotropic network model using the average simulated structure of (D) rat and (E) shrimp Tpm.

Domain structure in a rigid body model could be formed by border/hinge regions that with uncorrelated, local dynamics that interrupt dynamic communication between domains. A dynamic domain architecture with hinge regions is observed in mammalian Tpm, while crustacean Tpm produces exclusively local correlations (Figs. 4, **S3**). The covariance matrix derived from the rat and pig Tpm simulations suggests three major dynamic domains identified by hinge regions near residues 60 and 230 (Figs. 4B, **S3A-B**). The middle domain is the largest of the three domains. However, it is not uniform with long distance correlations at the C-terminal portion. Inter-domain correlations between the middle domain and both N- and C-terminal domains are also observed, with noticeable gaps in correlations across hinge regions.

Unlike mammalian Tpm, hinge regions are not observed in crustacean Tpm simulations (Fig. 4C). Dynamics are fundamentally local with even interchain correlations nonexistent (**Fig. S3C-D**). The narrow range of magnitudes in the correlation matrix correspond to the constrained fluctuations in the simulation. The rigid crustacean Tpm lacks clear hinge regions that could form dynamic domains.

The impact of the coiled-coil geometry alone on conformational dynamics was explored using elastic network model (ENM). ENM reduces structure to a series of nodes and springs (36). In our model nodes represented Cα positions from the average structure from the MD trajectory. For calculations, we used an anisotropic network model (ANM), which is subtype of ENM. The method assumes a universal force constant between residue nodes within a cutoff distance to produce a Hessian matrix (37). The covariance matrix was then determined by the top twenty modes from the Hessian. Comparing covariance matrices derived from rat Tpm MD and ANM show a similar three domain architecture. (Fig. 4B, 4D). However, in shrimp Tpm, covariance produced by the ANM not only follows the three-domain pattern in seen in the rat homolog, but also has almost an order of magnitude greater correlation, indicating that a straight coiled-coil conformation should exhibit even clearer domains if coiled-coil geometry was the primary factor (Fig. 4E). The difference in dynamics between rat and shrimp Tpm is dependent on sequence-specific determinants, rather than geometry alone.

### Low Frequency Correlations and Hinge Points

Dynamic domains exhibit collective low frequency motions. The time dependence of a dynamic process maybe be used to separate collective and local motions. In this regard, autocorrelation of protein molecular dynamics trajectories yields patterns of time correlations that can be distinguished based upon their damping behavior (38, 39). While picosecond dynamics of sidechain motions may be described well as damped oscillators by the Langevin equation, larger proteins exhibit complex dynamics, in part caused by overlaid low frequency oscillations in the autocorrelation (35, 39). Cα coordinates of corresponding residues on the two chains were averaged into a residue-specific midpoint that represents the coiled-coil interface. Autocorrelations were then calculated from the root square fluctuation (RSF) of each midpoint for both rat and shrimp Tpm (Fig. 5A, 5D). Similar to the covariance matrix, low frequency oscillations were observed in rat Tpm in three regions, suggesting shared long-range dynamics regions (Fig. 5C). In shrimp, autocorrelation produced overdamped decays, and in four regions there was overlaid a secondary extended decay (Fig. 5F). No clear low frequency oscillations were observed for shrimp Tpm for the timescales simulated.

**Figure 5.**
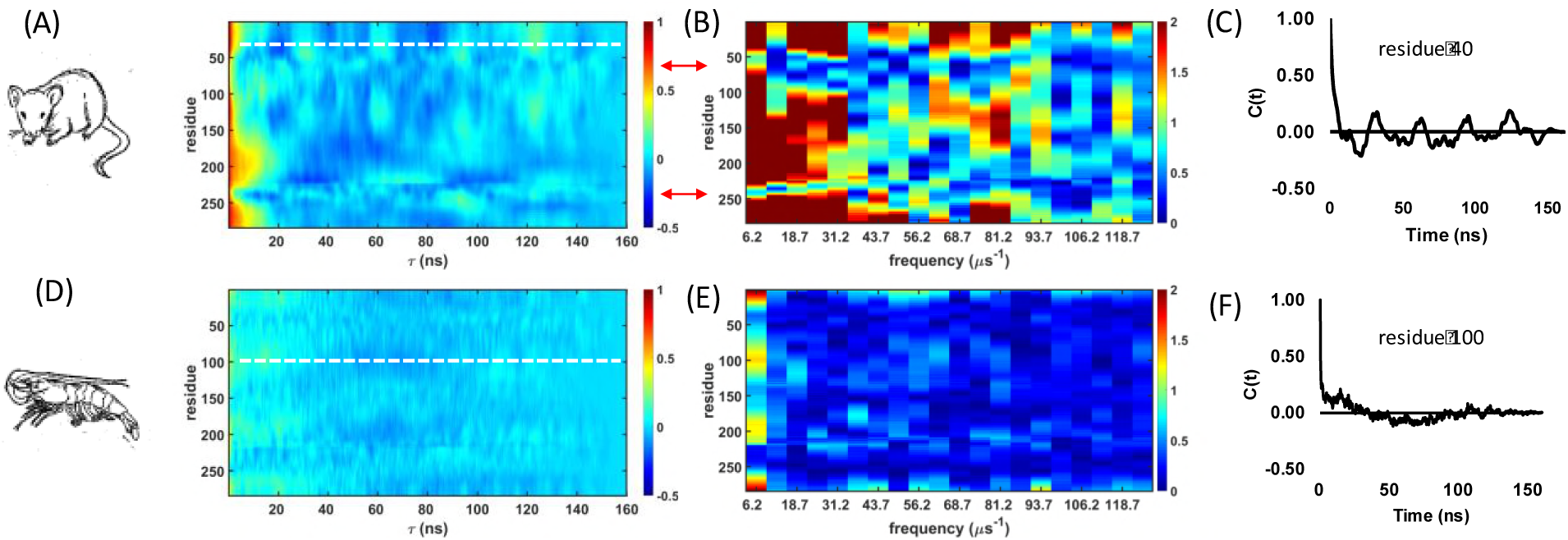
Time Series Autocorrelations. Autocorrelation functions were calculated from **(A)** rat and **(D)** shrimp MD using time series data of midpoint Cα RSF. Color bar indicates a normalized autocorrelation. **(B,E)** The fast Fourier transform of each residue autocorrelation was also computed, only the lowest 20 frequencies shown (right panel). Color bar represents the complex magnitude for each frequency. Red arrows highlight hinge points of higher frequency dynamics. **(C)** Correlation plot C(t) for rat Tpm, residue 40 shows low frequency undamped oscillations. Position shown as white line in (A). **(F)** In contrast, shrimp Tpm C(t) shows fast and slow damped oscillations. Position shown as white line in (D).

The power spectrum of RSF on a residue level provided complementary evidence of low frequency motions, particularly for vertebrate Tpms. The twenty lowest frequencies are shown, highlighting the differences noted prior in rat and shrimp Tpm. There are no identifiable shared low frequency correlations in shrimp or lobster Tpm (Fig. 5E, **S4**). Power spectra of rat and pig Tpm reveal three domains of shared low frequency amplitudes separated by high frequency borders (Fig. 5B). In the vertebrate Tpms, domains appear to form via low frequency dynamics. Higher frequency, local hinge regions form inter-domain boundaries. While three dynamic domains are identifiable in pig, the middle domain and C-terminal domain is less defined with fewer shared low frequency dynamics (**Fig. S4A**). In addition, the N-terminal border of the middle domain does not dampen low frequency motion as well as in the simulation with rat Tpm. Despite these differences between pig and rat, they behave differently to their crustacean counterparts in their ability to have shared low frequency dynamics across three domains.

### Mechanism of Dynamic Domain Formation

Curvature of the coiled-coil may be caused by deviations from the standard knob-into-holes packing in coiled-coils (4). Features of core residues, such as asymmetry due to small or bulky core residues or gaps in the interhelical interface, promote bending along the coiled-coil (40). The most well understood of these features is axial staggers between chain produced by alanine clusters in the core, which have been observed in multiple crystal structures(4, 16, 41). Axial shifts do not just locally distort structure, but correlations in the staggering direction between the nearest neighbors can induce long range curvature (42).

Rat and shrimp Tpm homologs largely share alanine cluster positions in the core, with the marked difference occurring by the absence of a second cluster between residues 74-81 (Fig. 6). Zheng et al. used metrics to calculate local interhelical dynamics in an MD simulation by measuring core residue distances (22). They found correlations between position of alanine clusters, decreased interhelical distance and large standard deviations of axial shifts. Similarly, our simulations for all four Tpms demonstrated a periodic tightening of coiled-coil packing that correspond to alanine clusters (**Fig. S5**).

**Figure 6.**
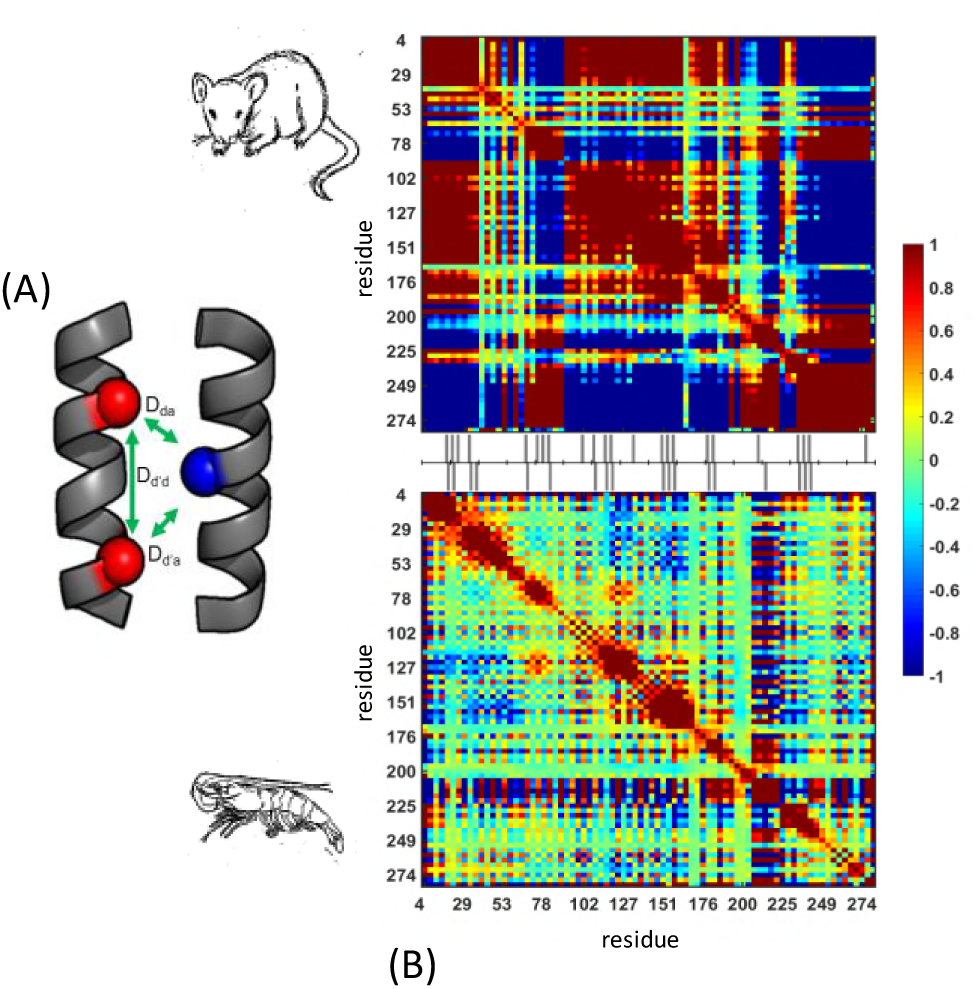
Domain mapping by correlations in coiled-coil axial shifts. (A) A schematic detailing calculation for axial shifts between core residues. The difference between the squared distance of each core Cα (blue) and its adjacent residues is calculated. For example, with respect to the a given ‘a’ position: 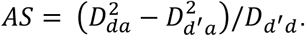
Axial shifts for core ‘d’ positions are calculated in a similar manner. (B) A covariance matrix of axial shift motions between core residues is derived from rat (top) and shrimp (bottom) Tpm MD trajectories. Color bar indicates negative and positive correlations. Lines adjacent to covariance matrix display position of core alanines for rat (top) and shrimp (bottom) respectively.

However, mean and standard deviations of axial shift corresponds less to core alanine clusters (Fig. 6C-D, **S5**), suggesting axial shifts to extend beyond the local dynamics of alanine clusters.

To visualize long-range correlations of axial shifts across the Tpm coiled-coil, we calculated a covariance matrix directly from axial shifts with respect to every core residue position (Fig. 6). Positive and negative covariance correspond to directional differences in shifts. In rat Tpm, four positively correlated off-diagonal blocks are observed. Off-diagonal blocks encompass and extend from alanine clusters. However, correlated axial shifts are interrupted at two border regions with near zero covariance. These border regions are near residues 60 and 220, which correspond to hinge points identified both by the covariance matrix and power spectrum from backbone Cα coordinates. The middle domain consists of two separate positively correlated blocks, which accounts for the asymmetry seen in previously stated analysis. In the pig Tpm MD simulations, core Ala clusters produce local shifts but no pattern in long-range correlations is observed (**Fig. S6**). As aforementioned, the pig simulation has less defined middle domain dynamics, which may account for its lack of long-range correlations. Importantly, despite sharing the majority of core alanines with rat, shrimp and lobster Tpm does not display any long-range off-diagonal blocks, in keeping with the essentially local and largely random nature of its dynamics (Figs. 6, **S6**).

## CONCLUSIONS

Long time scale molecular dynamics of Tpms uncovered distinct behaviors of mammalian and crustacean homologs. The dichotomy in dynamic behavior between crustacean and mammalian Tpm is perplexing due to the significant overlap in sequence and structure. Three dynamic domains in mammalian Tpms were derived from a long-range communication in core dynamics. Crustacean Tpms behaved as a rigid, straight rod dominated by local, high frequency dynamics.

Collective dynamics in mammalian Tpm are related to molecular hinges, which interrupt structural connectivity and allow for flexibility (43). While these regions produce a curved overall structure in keeping with the gestalt model, the coiled-coil retains flexibility with large scale motions between opposing dynamics through hinge regions (44). The middle dynamic domain itself may be flexible given that it consists of two different regions of correlated axial shifts. Taken together, the dynamic structure of mammalian Tpm provides for flexible conformational sampling that can adapt to F-actin binding.

Crustacean Tpm produces a rigid and linear structure without any long range dynamic correlations. By the gestalt model, its linear conformation would not be ideal for F-actin binding. The lack of correlations is due to the tight conformational space sampled, with an order of magnitude smaller fluctuations in RMSD and radius of gyration. The combined lack of large scale dynamics or the presence of hinge regions prevents the formation of dynamic domains. It may be possible that at longer conformational sampling shrimp Tpm has a curved conformation similar to the Lorenz-Holmes Tpm model. In terms of the dynamics the rigid global structure of crustacean Tpm is likely to adopt a preset, stable conformation for interacting with F-actin.

It is perhaps not surprising that salient features of tropomyosin sequence and structure have remained unchanged over one half billion years of evolution. Actin participates in protein-protein interactions with hundreds of other partners (45), which would be expected to significantly constrain its rate of evolution (46). This then constrains the evolution of actin binding sites and structural evolution of Tpms. However, adaptations of Tpm dynamics in response to temperature have been observed (47, 48), highlighting the importance of considering the impact of protein biophysical properties on evolution (49). As molecular dynamics simulations become computationally accessible, it becomes increasingly feasible to perform comparative long time-scale simulations to examine the role of protein dynamics in shaping molecular adaptation and divergence (50, 51).

## METHODS

### Molecular Dynamics

The 7 Å crystal structure of the pig Tpm was used (PDB ID: 1C1G) as the initial structure for MD simulations (27). The rat, shrimp, and lobster homologs were created by changing the amino acid sequence using the protCAD (protein Computer Aided Design) software (29). The rat Tpm sequence is from *Rattus norvegicus* alpha-1 chain isoform (gi 671696014); the pig homolog from *Sus scrofa* alpha-1 chain isoform (gi 148222268); shrimp homolog from *Penaeus monodon* (gi 60892782); lobster homolog from *Homarus americanus* fast isoform (gi 2660868).

Molecular dynamics simulations were performed using AMBER14 (26). The ff14sb force field was used for molecular mechanics. The protein was surrounded with TIP3P water box over a radius of 10 Å from the protein. Net charge was set to zero by adding Na+ and Cl- to the system. Initially, two rounds of optimization were performed to energetically minimize the structure, starting with the steepest descent and then the conjugate gradient method. MD calculations wer performed starting with a 2 fs time-step required for the SHAKE algorithm to constrain bond lengths. Periodic boundaries were set under constant volume. The system was thermalized from 0 to 300K using Langevin dynamics using a collision frequency of 3 ps^−1^. Next, equilibration was performed under constant temperature, no pressure scaling for 100 ps. The final MD trajectory was run with coordinates written to trajectory file every 20 ps. Trajectory coordinates were analyzed with pytraj, a CPPTRAJ based python module (52).

### Anisotropic Network Model

An average structure of each MD trajectory was created using CPPTRAJ. The ANM was created with Cα from average structure using the ProDy package (53). The force constant for ANM was set as 1.0 kcal/Å^2^ and a cut-off distance 13.5 Å as constraints for interactions based on previously derived general ranges for proteins (37). The program created a Hessian matrix, which was diagonalized. The top twenty eigenvectors were used to create the covariance matrix for Cα fluctuations.

### MD analysis

All MD analysis was done using the MATLAB software package (54). Distribution of conformational space by radius of gyration and RMSD using a kernel density smoothing function (ksdensity). PCA was calculated from a covariance matrix derived from Cα fluctuations relative to the mean position using the eigen-decomposition function (eig). The root square fluctuation of Cα position was used to calculate autocorrelation (autocorr) were transformed to the frequency domain with a fast Fourier transform (fft). Sequence analysis was done using a Needleman-Wunsch alignment algorithm (nwalign).

## SUPPORTING MATERIAL

Figures S1–S6 and Table S1 included in the supporting material.

## AUTHOR CONTRIBUTIONS

JKJ performed the experiments. JKJ and VN designed the experiments, analyzed the data and wrote the paper.

## ACKNOWLEDGEMENTS

We thank Sarah Hitchcock-DeGregori and Douglas Pike for useful scientific discussions. This work was supported by the National Institutes of Health R21 AI-088627.

